# Bayes Estimate of Primary Threshold in Cluster-wise fMRI Inferences

**DOI:** 10.1101/2020.10.03.324962

**Authors:** Yunjiang Ge, Stephanie Hare, Gang Chen, James Waltz, Peter Kochunov, L. Elliot Hong, Shuo Chen

## Abstract

Cluster-wise statistical inference is the most widely used technique for functional magnetic resonance imaging (fMRI) data analyses. Cluster-wise statistical inference consists of two steps: i) primary thresholding that excludes less significant voxels by a pre-specified cut-off (e.g., *p* < 0.001); and ii) cluster-wise thresholding that controls the family-wise error rate (FWER) caused by clusters consisting of false positive suprathreshold voxels. It has been well known that the selection of the primary threshold is critical because it determines both statistical power and false discovery rate. However, in most existing statistical packages, the primary threshold is selected based on prior knowledge (e.g., *p* < 0.001) without taking into account the information in the data. In this manuscript, we propose a data-driven approach to objectively select the optimal primary threshold based on an empirical Bayes framework. We evaluate the proposed model using extensive simulation studies and an fMRI data example. The results show that our method can effectively increase statistical power while effectively controlling the false discovery rate.

## 1 Introduction

Functional magnetic resonance imaging (fMRI) technique has become a popular tool for non-invasively studying circuit-level brain activity for more than two decades. Statistical analyses of neuroimaging data remain challenging due to their high-dimensionality and spatiotemporal dependence structure (Derado et al., 2010; Lindquist, 2020; Lindquist & Mejia, 2015; Smith & Nichols, 2018). Advanced statistical methods have been developed to solve the multiple comparison problem and successfully applied to many fMRI studies (Alberton et al., 2020; Chen et al., 2019; Nichols, 2012; Smith & Nichols, 2009). Among these statistical methods, the cluster-wise inference remains the most commonly used tool for neuroimaging data analysis due to its relatively high sensitivity and low computational cost comparing to voxel-extent based thresholding methods (Friston et al., 1994; Woo et al., 2014). The performance and parameters of this procedure have been well discussed and studied (Eklund et al., 2018; Eklund et al., 2016; Flandin & Friston, 2019; Y.-W. Hong et al., 2019).

### A spatial point process perspective of cluster-wise inference

Cluster-wise inference consists of two steps: a primary thresholding step that applies a cut-off to all voxels and only keeps those supra-threshold voxels; and a cluster-extent-based thresholding step to avoid selecting false positive clusters under the null hypothesis (e.g., no activation). In this current study, we focus on the non-parametric inference method, due to its robustness although parametric methods (e.g., random field theory) may be more efficient when the assumptions are well met (Bennett, Wolford, et al., 2009; Eklund et al., 2015; Hayasaka & Nichols, 2003; Nichols, 2012; Schwartzman & Telschow, 2019). The high sensitivity of cluster-wise inference comparing to other multiple-testing correction methods (e.g., false discovery rate - FDR control) is built on the appropriate modeling of the stochastic spatial process for informative voxels. Specifically, the primary thresholding in step one binarizes all voxels to ones and zeros, and then the statistical inferences for the whole brain become a spatial point process in a 3D brain space where only supra-threshold voxels present as points (Kang et al., 2011). Under the null hypothesis, no brain area is associated with the external covariates, and the false positive points (voxels false-positively surviving the primary threshold) are assumed to follow a homogeneous spatial process. Under the alternative hypothesis, the identification of true events (voxels true-positively surviving the primary threshold) is a non-homogeneous/clustered process. Asymptotically, the combinatorial probability that true events connect (spatially adjacent) to each other and form a large cluster is much greater than the combinatorial probability of false positive voxels connecting to each other and forming a cluster with an equal or greater size. In that, we assume the noise is not related to any brain function or brain structure. However, the calculation of the asymptotic combinatorial probability is difficult because the relationship of the covariance between voxels and their geometric distance can be non-stationary and non-linear (Cressie, 2015). Fortunately, nonparametric test methods (e.g., permutation tests) can provide a good approximation of the asymptotic combinatorial probability of false positive voxels forming a cluster with a certain size since the empirical covariance between voxels is preserved in each permutation (Eklund et al., 2016; Nichols, 2012; Nichols & Hayasaka, 2003). Nonparametric tests are also widely and successfully applied in other spatial point models, for example, SaTscan (Kulldorff, 2006) is used to monitor/detect a clustered spatial point incidences (disease outbreaks). Therefore, cluster-wise inference gains additional sensitivity by capitalizing on the patterns of positive voxels in a non-homogeneous process. However, the price for the additional power is the loss of local power (i.e., without voxel-level inference).

A key limitation for cluster-wise inference is its venerability to the poor selection of the voxel-level primary threshold. An overly-conservative threshold may lead to trivial clusters of connected true positive voxels and cause low statistical power. On the other hand, a liberal primary threshold (e.g., *p* < 0.01) can generate massive false positive points in a smoothed brain space and thus can be connected and form false positive clusters (Woo et al., 2014). Both false positive clusters and low statistical power become major potential causes of the low reproducibility and replicability of fMRI findings (Lindquist, 2020).

Setting the primary threshold at *p* < 0.001 is now standard for most studies, because it can generally effectively control false positive findings based on empirical studies (Bennett, Miller, et al., 2009; Woo et al., 2014). However, a pre-specified voxel-wise threshold may be sub-optimal because it does not account for several important factors of the data, including sample size, effect size, noise level, and the selection of statistical models, among many others. We illustrate the concept of data-driven optimal threshold selection by a toy example (Figure 1). We consider a simple scenario that all voxels in a brain image can be divided into two sets: those are truly associated with the covariate of interest and the rest. The test statistics of the two sets follow a non-null distribution and a null distribution, respectively. We argue that the optimal primary threshold should be selected based on the non-null and null distributions rather than a pre-specified threshold. For example, if the two distributions are well separated (large sample sizes or significantly strong signals), a more rigorous primary threshold (e.g., more stringent than 0.001) should be used, to suppress the false positive findings. In contrast, if the two distributions are less separable (but still separable, exhibiting small/moderate sample sizes and moderate/large effect sizes), a less conservative primary threshold (e.g., less conservative than 0.001) should be used to keep the false discovery rate at a low level while maintaining a maximal statistical power. Therefore, empirical distributions of null and non-null distribution can reflect the sample size, effect size, and noise level from the data and provide important guidance for primary threshold selection.

**Figure 1:**
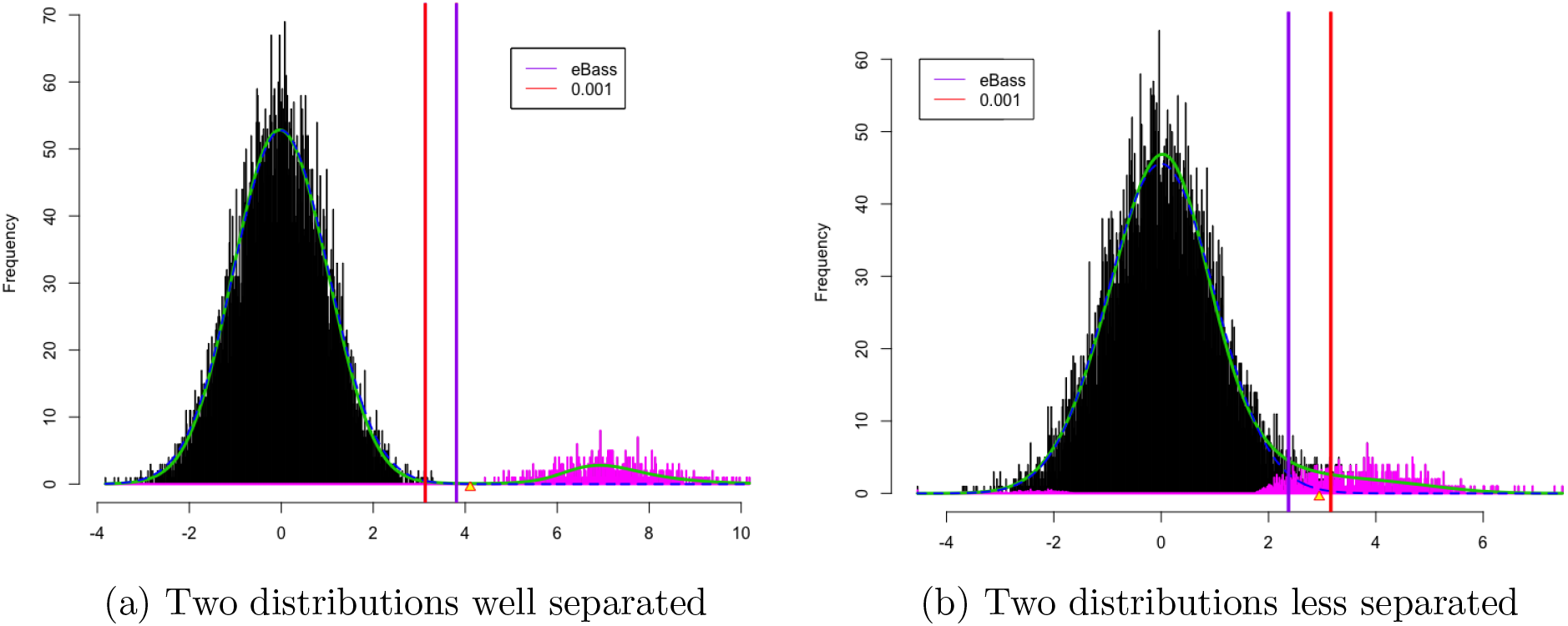
Non-null (in purple) and null (in black) distribution of test statistics from the (non-)event spatial points. The red vertical line marks down the corresponding z-score when the p-value is 0.001 on each graph. Yellow triangles on the x-axis indicate threshold z-values for local fdr 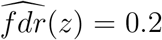 if such cases exist. The null sub-density *π*_0_*f*_0_ is marked in a blue dashed line, and the mixture density is marked in a solid green line.

In the above example, we show that a data-driven primary threshold can maximize the statistical power to detect true positive clusters while effectively controlling the false discovery rate. However, the data-driven primary threshold selection procedure has not been fully developed. To fill this gap, we propose a new **e**mpirical **B**ayes **A**daptive Thre**s**hold **S**election (eBass) method to objectively select the optimal primary threshold selection based on the information from data. The eBass objective function aims to achieve a maximal statistical power while avoiding false positive voxels connected to be a false positive cluster. We develop new algorithms to implement the objective function and provide the corresponding theoretical properties. In this paper, we focus on the two-step cluster-wise fMRI inference with nonparametric statistical tests by providing a new primary threshold selection strategy. We note that alternative advanced statistical methods, including both nonparametric inference methods (e.g., threshold-free methods cluster enhancement - TFCE and pTFCE (Smith & Nichols, 2009; Spisák et al., 2019)) and parametric inference (e.g., frequentist and Bayesian models (Benjamini & Heller, 2007; Chen et al., 2019; Schwartzman & Telschow, 2019)), can produce reliable and biologically meaningful results. Thus, eBass can become a complement to these commonly-used statistical approaches and enhance the widely used two-step cluster-wise inference.

The rest of this paper is organized as follows: 1) we introduce eBass method and algorithm in section 2; 2) we perform extensive simulation analysis to fully assess the properties of eBass, in Section 3; 3) we apply eBass to an fMRI data analysis for schizophrenia research and conclude with discussions and future works.

## 2 Method

We consider the multiple comparison problem for all brain voxels in a spatially smooth 3D space. For example, the voxel-level brain activation to an external stimulus (task-induced fMRI) and seed voxel based brain connectivity map (resting-state fMRI). Let *v* be an index for brain voxel and *v* = 1,⋯ , *V*. We perform statistical inference on each voxel marginally (e.g., by a general linear model GLM), and conditionally (Bowman, 2005; Derado et al., 2010; Risk et al., 2016). Therefore, there are simultaneously *V* hypothesis tests with their test statistics ***Z*** and corresponding *p* values **P**:

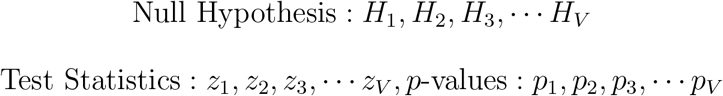

The commonly-used multiple testing correction methods (Benjamini-Hochberg false discovery rate, or BH-FDR, correction) correct the multiplicity at the voxel level. The two-step cluster-wise inference aim to extract cluster-level findings and gain additional power (Nichols & Hayasaka, 2003). Specifically,

1. We consider the primary thresholding as a screening step. We first apply a pre-determined threshold *θ* to binarize all voxels based on their *p*-values. Denote the indicator variable *δ_v_ = I*(*p_v_ < θ_p_*) (e.g., *θ_p_* < 0.001) and Δ = {*δ_v_* > 0}, where *I* is an indicator function. The binarization naturally leads to voxel-level false positive rate and sensitivity.
2. Perform permutation tests using the cluster extent as a test statistic to select a set of spatially adjacent supra-threshold voxels Δ as the resulting cluster while controlling the family-wise error rate (FWER). This step bears a resemblance to the commonly used spatial statistical models (e.g., SaTScan) that can competently handle an inhomogeneous spatial point process with clustered patterns (Waller & Gotway, 2004). Thus, the final output is cluster-level findings (adjusting FWER).

We refer voxel-level false discovery rate as 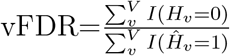, and cluster-level family-wise error as cFWER=Pr(at least one detected cluster is false positive). The two-step clusterwise inference controls cFWER instead of vFDR. Nevertheless, the vFDR can be used to evaluate the selected primary threshold in step one (screening).

It has been well-known that the choice of primary threshold is critical because i) an over conservative primary threshold can achieve a low voxel-level false discovery rate at the cost of low sensitivity (the cluster-level sensitivity is also low because true positive voxels are few to form clusters); ii) a liberal primary threshold can lead to detecting both true and false positive clusters (i.e., high cFWER). Either low sensitivity or high cFWER can lead to less replicable results because i) the probability is low to observe overlapped true findings across data sets with low sensitivity, and ii) the chance for false positive voxels reappearing in different data sets is small. Currently, the primary threshold of *p* < 0.001 is well accepted by the research community, while *p* < 0.01 is considered overly-liberal (Eklund et al., 2018). Here, we argue that a data-driven primary threshold may better balance the above trade-off than the pre-determined primary threshold.

We aim to select an optimal primary threshold to achieve maximal sensitivity (power) with a low false discovery rate at the voxel-level. In practice, however, neither voxel-level sensitivity nor FDR is known because the ground-truth is unavailable. We resort to an empirical Bayes framework for calculating the estimated voxel-level sensitivity and FDR.

### 2.1 Empirical Bayes estimated voxel-level sensitivity and FDR

The empirical Bayes framework has been developed to estimate the marginal distribution of the null and non-null test statistics for the multiple testing problem (Efron, 2012; Fan et al., 2012). In these models, the test statistics of the whole brain voxels follow a mixture distribution, *f*(*z*) = *π*_0_*f*_0_(*z*) + *π*_1_ *f*_1_(*z*), where *π*_0_, *π*_1_(*π*_1_ = 1 − *π*_0_) are the prior probabilities for a voxel belonging to the null and non-null components.

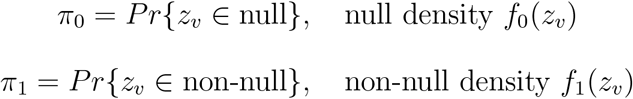

The *posterior* probability of a voxel is from the null set given **z** is Pr{*z_v_* ∈ null|**z**} = *π*_0_*f*_0_(*z*)/*f*(*z*). A critical step of the empirical Bayes method is to estimate the mixture density 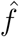. Fortunately, numerous efficient and robust numerical algorithms (e.g., MLE based and Poisson regression estimates) have been developed (Efron, 2014). The null density 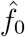 can be estimated by maximum likelihood estimation and central matching method. The prior probability *π*_0_ is estimated based no the estimated null distribution, and accordingly, 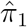 is 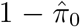. Generally, the empirical Bayes estimation method provides consistent and robust estimates for 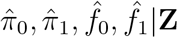 based on data **Z** (Efron, 2012; Petrone et al., 2014).

Given estimated Pr{*z_v_* ∈ null} at each voxel, we can calculate the *posterior* sensitivity and FDR across all voxels at a cut-off *z_θ_* by:

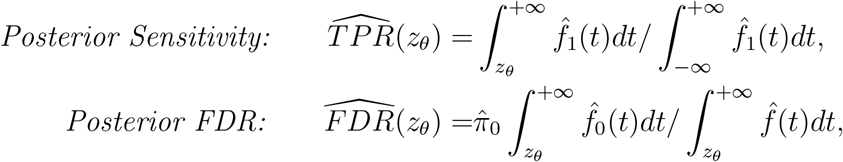

and accordingly the true discovery rate (TDR) is

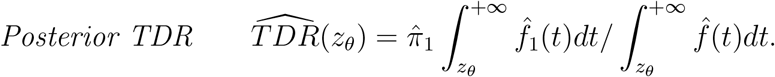

The 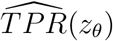 and 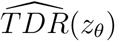 inherit the consistency property from empirical Bayes estimators (see Theorem 2 with detailed proofs in Appendix). Therefore, the empirical Bayes estimated 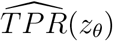 and 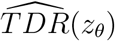 provide satisfactory surrogates for the true, yet unknown, sensitivity and FDR, which are required to determine the optimal threshold.

### 2.2 Objective function for the optimal threshold

Built on the EB *posterior* voxel-level sensitivity and FDR (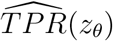 and 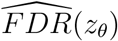), we propose an objective function for optimal threshold selection. Specifically,

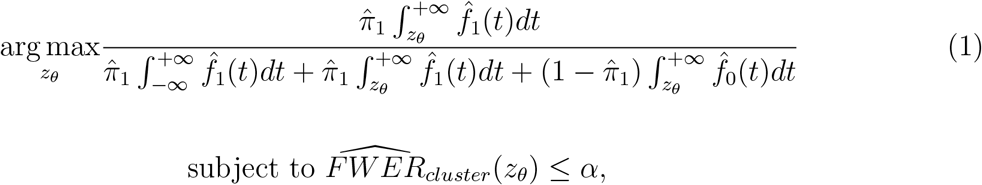

where the optimal cut-off *z_θ_* is the estimand. The objective function (1) is the harmonic mean of empirical Bayes sensitivity and true discovery rate, and provides the optimal selection of 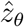 that maximizes the power with the family-wise error rate rate controlled under the level of *α* (see details of Theorem 1 in Appendix). The objective function is expressed by the empirical Bayes *posterior* parameters and functions 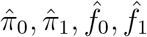.

The cluster level family-wise error rate 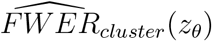 is determined by the number of estimated false positive voxels 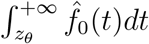 and the cluster size cut-off. Specifically, we denote the estimated number of false positive voxels using a cut-off *z_θ_* by 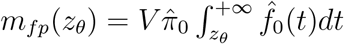. Next, we implement the permutation procedure by relabelling to determine the clustersize cut-off with the FWER level *α* and primary threshold *z_θ_*. Let *K_α_*(*z_θ_*) be the permutation determined cluster-size cut-off. When *z_θ_* is liberal, *m_fp_*(*z_θ_*) tends to be a large number and thus is more likely to form large clusters composed by false positive voxels surpassing the cut-off *K_α_*(*z_θ_*). In contrast, a stringent *z_θ_* leads to a small *m_fp_*(*z_θ_*) prohibiting false positive voxels connecting into clusters greater than *K_α_*(*z_θ_*). 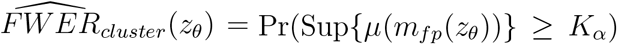, where *μ* is the cardinality measure of any set of contiguous voxels formed by *m_fp_*(*z_θ_*) in the brain space. Then, we define the search domain for 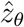 by Ω_*α*_ = {*z_θ_*: Pr(Sup{*μ*(*m_fp_*(*z_θ_*))} ≥ *K_α_*) < *α*}. We provide the computational details to calculate Sup{*μ*(*m_fp_*(*z_θ_*))} estimate Ω in the Appendix. Next, we optimize *z_θ_* on the support Ω.

We summarize the computational procedure of eBass in three steps:

Step 1: Calculate 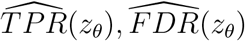 with the empirical Bayes estimated 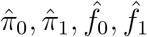, and the objective function as

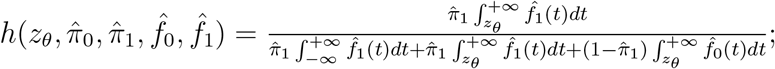
Step 2: Identify the support Ω_*α*_ which guarantees 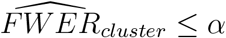 (see Appendix).
Step 3: Optimize the *z_θ_* by 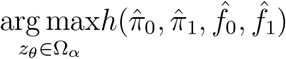 subject to constraint in Step 2, under the regularity condition for Cost-Sensitive Classification algorithm (Eban et al., 2017). According to the algorithm, the objective function *h*(·) reduces to a weighted classification problem, where the errors are approximated by cost-sensitive binary classification algorithms (e.g., logistic regression) with asymmetric costs.

The optimization of (1) yields the optimal primary threshold 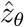, which maximizes the sensitivity with 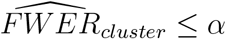. Note that 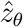 is objectively selected by fully leveraging the information from the empirical data. In practice, the sample size, effect size, noise level, spatial dependence across all voxels of the empirical data set can influence the separability of the null and non-null distribution, and thus the optimal primary threshold should be updated to accommodate these factors. The empirical Bayes estimated FDR and sensitivity determines our objective function (1), and bridges the data information and parameter optimization. The data-driven primary threshold selection method eBass automatically maximizes the empirical Bayes sensitivity while controlling voxel-level FDR and then cluster-level FWER. Therefore, eBass primary threshold can outperform the pre-specified primary thresholds in many scenarios (see simulation and data example results).

In summary, we provide a data-driven optimal primary threshold selection step via an empirical Bayes framework. The selection of the primary threshold is more flexible because eBass optimizes the primary threshold by maximizing the sensitivity while rigorously controlling the cluster-wise false positive error rate. We provide theoretical proof details for the optimality (Theorem 1 in Appendix) and consistency (Theorem 2 in Appendix). In the following simulation analysis and real data example, we demonstrate that eBass can improve the statistical power without losing the rigor of FWER.

## 3 Simulation

### 3.1 Data Description

In the simulation study, we evaluate the performance of eBass and compare it to the existing methods. We first simulate 2D image data for multiple subjects. The number of voxels in each image is *V* = 100 × 100 = 10, 000, and thus the number of simultaneous tests is 10,000. We assume that most voxels are from the null set, whereas two squared areas (*N*_0_ = 21 × 21 + 6 × 6 = 477 voxels) are from the non-null, see Fig 2b. We apply a commonly-used two-group (i.e., cases vs. controls) scenario, which can be easily extended to the regression setting. First, let voxels from the null set follow a normal distribution *N*(0, 1) for both cases and controls. Within the two squared areas, the non-null voxels of the cases follow a normal distribution *N*(*μ_k_*, 1) (*k* = 1, 2 for the two areas), whereas the voxels of the controls follow a *N*(0, 1) distribution. The signal-to-noise ratio (SNR) as the reciprocal of the coefficient of variation, *SNR* = *μ_k_/σ*, the *σ* = 1 allows the difference of group means to be the true positive effect size (ES) which is equivalent to Cohen’s d. A higher SNR can lead to higher sensitivity and a lower FDR, and vice versa. We further smooth each image with a Gaussian filter, using a full width at half maximum (FWHM) equivalent to 8mm. The voxels in the smoothed image are correlated like the real fMRI data. We further let the number of subjects per group be 30, 60, and 100. For each setting, we simulate 100 data sets.

**Figure 2:**
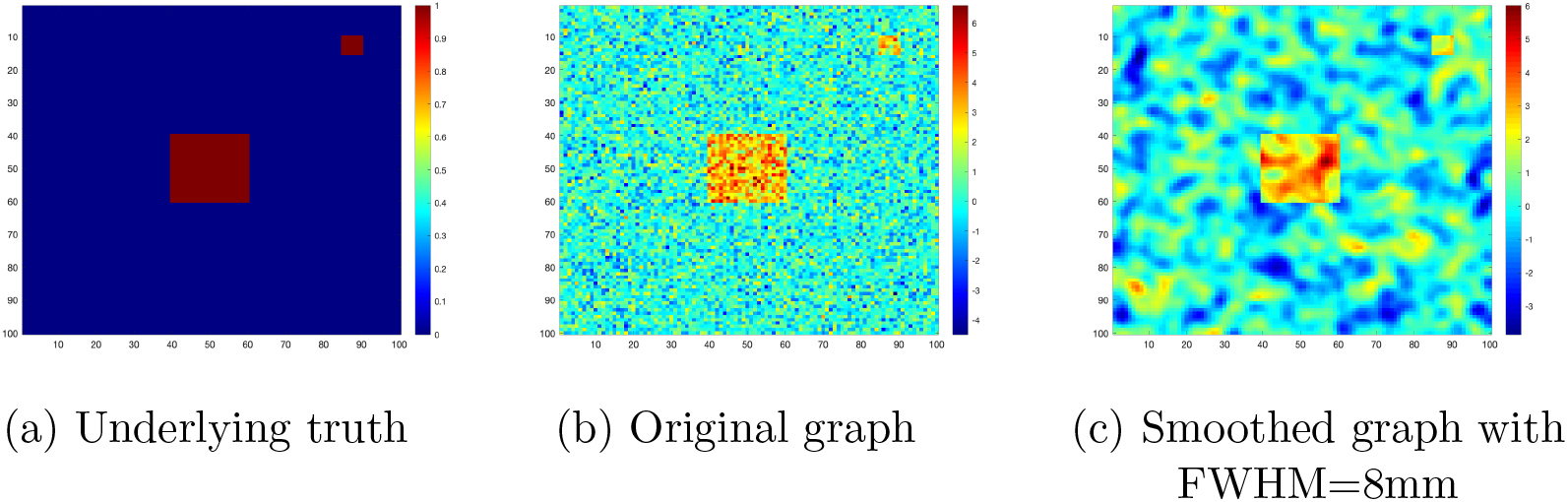
(a) is the graph of the underlying truth. Red squares represent the activated regions. (b) shows one original graph with sample size 30 per arm, an effect size set to 0.8. Non-activated voxels in blue region of (a) generated that follow *N*(0, 1) and activated voxels in red regions of (a) follow *N*(0.8, 1). (c) was the original graph in (b) smoothed with a Gaussian kernel of FWHM=8mm.

### 3.2 Data analysis

For each data set, we perform the two-step cluster-wise inference. We determine the primary threshold using eBass as well as a variety of popular methods, including BH-FDR correction, *p* < 0.001, and *p* < 0.01. We evaluate the performance of these methods in terms of voxel-wise sensitivity (Sensitivity) and FDR (vFDR), together with cluster-wise FWER (cFWER) by comparing the resulting clusters selected by the cluster-wise inference to the two true squares. Note that ultimately, the voxel-wise sensitivity and FDR are not applicable in cluster-wise inference, we yet provide them in order to evaluate the quality of controls in the voxel-level primary thresholding step.

### 3.3 Results

We summarize simulation results in Table 1. We first compare the results when the effect size is medium (ES=0.6). For the sample size of 30 cases vs. 30 controls, the study is underpowered and the test statistics from the true positive voxels are mixed with false positive voxels (Cremers et al., 2017; Lindquist et al., 2008). Therefore, only a few voxels can survive the corrected primary threshold and form a cluster with a size greater than the step two cluster-level threshold. The vFDRs for all methods are well controlled. The Sensitivity of eBass is 137% higher that *p* < 0.001 threshold and 388% higher than BH-FDR correction. Although the well-controlled vFDR prohibits false positive findings, the overall low sensitivity can also lead to a low replicability because true positive findings are rarely overlapped across data sets.

**Table 1.** Simulation result for original images of ES=0.6, 0.8 and 1.0, sample size 30, 60 and 100 per arm smoothed with FWHM=8mm

| | | eBass | BH-FDR | $p < 0.001$ | $p < 0.01$ |
| --- | --- | --- | --- | --- | --- |
| Effect Size = 0.6 |  |  |  |  |  |
| Sample size<br>30 per arm | Threshold | 0.0027(Q1)/0.0051(Q2)/0.0147(Q3) | 0.0002(Q1)/0.0009(Q2)/0.0017(Q3) | 0.001 | 0.01 |
|  | Sensitivity | 0.1932±0.0934 | 0.0396±0.0325 | 0.0816±0.0344 | 0.2263±0.081 |
|  | vFDR | 0.024±0.0653 | 0 | 0.0117±0.0469 | 0 |
|  | cFWER | 5.6% | 0 | 6.3% | 0 |
| Sample size<br>60 per arm | Threshold | 0.0012(Q1)/0.0027(Q2)/0.0056(Q3) | 0.0008(Q1)/0.0009(Q2)/0.0013(Q3) | 0.001 | 0.01 |
|  | Sensitivity | 0.5039±0.1237 | 0.3731±0.1481 | 0.3777±0.1075 | 0.6763±0.1164 |
|  | vFDR | 0.0067±0.0155 | 0.0050±0.0154 | 0.0046±0.0144 | 0.0127±0.0256 |
|  | cFWER | 5% | 10% | 10% | 10% |
| Sample size<br>100 per arm | Threshold | 0.0009(Q1)/0.0020(Q2)/0.0032(Q3) | 0.0020(Q1)/0.0021(Q2)/0.0022(Q3) | 0.001 | 0.01 |
|  | Sensitivity | 0.8292±0.0749 | 0.8460±0.0767 | 0.7850±0.081 | 0.9378±0.0422 |
|  | vFDR | 0 | 0.0019±0.0049 | 0.0001±0.0006 | 0 |
|  | cFWER | 0 | 0 | 0 | 0 |
| Effect Size = 0.8 |  |  |  |  |  |
| Sample size<br>30 per arm | Threshold | 0.0073(Q1)/0.0097(Q2)/0.0106(Q3) | 0.0006(Q1)/0.0007(Q2)/0.0009(Q3) | 0.001 | 0.01 |
|  | Sensitivity | 0.6012±0.0886 | 0.2881±0.1141 | 0.3146±0.0848 | 0.6109±0.0914 |
|  | vFDR | 0.004±0.01 | 0.0045±0.0201 | 0 | 0.0039±0.0096 |
|  | cFWER | 0 | 5% | 0 | 0 |
| Sample size<br>60 per arm | Threshold | 0.0034(Q1)/0.0050(Q2)/0.0055(Q3) | 0.0021(Q1)/0.0023(Q2)/0.0023(Q3) | 0.001 | 0.01 |
|  | Sensitivity | 0.9438±0.0396 | 0.9072±0.0552 | 0.8559±0.0605 | 0.9624±0.0376 |
|  | vFDR | 0.0109±0.0252 | 0.0052±0.0156 | 0.0049±0.0129 | 0.0119±0.0263 |
|  | cFWER | 15% | 10% | 15% | 10% |
| Sample size<br>100 per arm | Threshold | 0.0011(Q1)/0.0016(Q2)/0.0020(Q3) | 0.0024(Q1)/0.0024(Q2)/0.0025(Q3) | 0.001 | 0.01 |
|  | Sensitivity | 0.9945±0.0095 | 0.9955±0.0087 | 0.9897±0.0145 | 0.9988±0.0030 |
|  | vFDR | 0.0036±0.0095 | 0.0023±0.0073 | 0.0025±0.0078 | 0.0064±0.0147 |
|  | cFWER | 10% | 5% | 10% | 5% |
| Effect Size = 1.0 |  |  |  |  |  |
| Sample size<br>30 per arm | Threshold | 0.0093(Q1)/0.0119(Q2)/0.0154(Q3) | 0.0014(Q1)/0.0016(Q2)/0.0018(Q3) | 0.001 | 0.01 |
|  | Sensitivity | 0.8539±0.0522 | 0.6306±0.1122 | 0.5781±0.0982 | 0.8432±0.0526 |
|  | vFDR | 0.0093±0.0189 | 0.0045±0.0138 | 0.0026±0.0115 | 0.0057±0.0168 |
|  | cFWER | 5% | 10% | 5% | 5% |
| Sample size<br>60 per arm | Threshold | 0.0015(Q1)/0.0022(Q2)/0.0031(Q3) | 0.0023(Q1)/0.0024(Q2)/0.0025(Q3) | 0.001 | 0.01 |
|  | Sensitivity | 0.9851±0.0154 | 0.9842±0.0191 | 0.9727±0.0281 | 0.9951±0.0087 |
|  | vFDR | 0.0003±0.0014 | 0.0002±0.0006 | 0.0027±0.0085 | 0.0097±0.0177 |
|  | cFWER | 0 | 0 | 10% | 5% |
| Sample size<br>100 per arm | Threshold | 0.0004(Q1)/0.0006(Q2)/0.0008(Q3) | 0.0024(Q1)/0.0024(Q2)/0.0025(Q3) | 0.001 | 0.01 |
|  | Sensitivity | 1 | 1 | 1 | 1 |
|  | vFDR | 0 | 0.0015±0.0034 | 0 | 0.0067±0.0105 |
|  | cFWER | 0 | 0 | 0 | 0 |

When the sample size is increased to 60 subjects per arm, the statistical power increases to above 80% at each voxel. The vFDRs of eBass, BH-FDR correction, and *p* < 0.001 primary threshold are around 0.005, while vFDR of threshold *p* < 0.01 is close to 0.013. Note that our eBass primary threshold varies across repeatedly simulated data sets, and automatically adapt to the characteristics of the data. In that, our average vFDR is about the same as *p* < 0.001 and BH-FDR, but has a significantly increased Sensitivity. Since the sample size of 60 vs. 60 and ES=0.6 is common in fMRI studies, these results provide practical guidance for optimal primary threshold selection for cluster-wise inference.

Last, for sample size of 100 vs. 100, the test statistics of voxels from the null set are clearly apart from those from the non-null set, which leads to generally increased Sensitivity in all methods. The eBass method has slightly higher Sensitivity than that of *p* < 0.001 with better controlled vFDR because of its adaptive optimal threshold selection. The cFWER for all methods are well-controlled.

The results of larger effect sizes (ES=0.8 and 1) follow a similar pattern as above (see Table 1). When the sample size is small (i.e., 30 cases vs. 30 controls), eBass significantly improves Sensitivity while keeping the vFDR at a very low level. At effect size 0.8 and the sample size is medium to large (i.e., 60, 100 subjects per arm), adaptive threshold selection methods eBass and BH-FDR correction performed slightly superior to the fixed primary threshold *p* < 0.001. When the effect size reaches to 1.0, there is not much difference in Sensitivity among all methods especially when sample size is very large (100 subjects per arm). All methods control the vFDR well when effect sizes increase. Since we focus on cluster-wise inference and cluster-level FWER, we only compare eBass with existing cluster-wise inference methods. In the appendix, we show the comparison results of TFCE and cluster-wise inference methods including eBass by assuming all voxels in significant clusters as positive.

In summary, eBass shows advantageous performance in improving the sensitivity while controlling the voxel-wise FDR, specially when the sample size and effect size is small to medium. With increased sample size and effect size, most of the widely used thresholding methods tend to have a good performance with similar primary thresholds.

## 4 Data Example

### 4.1 Data Acquisition

Resting-state fMRI (Rs-fMRI) data were collected from 96 schizophrenia patients (SZs) at the University of Maryland Center for Brain Imaging Research. The average age of the SZ cohort is 35.9±13.2, 28 of the participants are females. A Siemens 3T TRIO MRI (Erlangen, Germany) system equipped with a 32-channel phase array head coil was used to collect the resting-state *T*2*-weighted images with the following parameters: TR=2s, TE=30ms, flip angle=90°, FOV=248mm, 128 × 128 matrix, 1.94 × 1.94 in-plane resolution, 4mm slice thickness, 37 axial slices, 444 volumes). During the scan, participants were asked to keep their eyes closed and relax.

### 4.2 Data Preprocessing

Pre-processing of the rs-fMRI data was performed using the Data Processing & Analysis for (resting-state) Brain Imaging (DPABI) toolbox (Yan et al., 2016). The first ten-time frames were removed to allow for signal stabilization. Raw data underwent motion correction to the first image, slice-timing correction to the middle slice, and normalization to MNI space. To ensure that spurious motion and physiological artifacts did not drive observed effects in our statistical analyses, resting data also underwent regression of 6-motion parameters and their derivatives (12 total motion estimates) and physiological (white matter and cerebrospinal fluid) signals prior to spatial smoothing with an 8mm FWHM Gaussian kernel. Framewise displacement was calculated for each image; this measure differentiates head realignment parameters across frames and generates a 6-dimensional times series that represents instantaneous head motion (Power et al., 2012). All individuals in the current analysis had mean framewise displacement ≤ 0.25 to better control for potential confounding effects of motion and motion artifacts on the rs-fMRI signal.

### 4.3 Data Analysis

We aimed to examine the association of functional connectivity of the default mode network with smoking status. The seed voxel method was used by placing a 10mm spherical seed centered on the precuneus/posterior cingulate centered (PCC). The correlations were calculated between the rest of voxels and the seed, and then normalized by the Fisher’s Z transformation. The two-step cluster-wise inference was conducted to identify smoking-related voxel-clusters.

In step one, voxel-wise regression was performed with the factor of interest (e.g., smoking status) on 215,348 nonzero voxels while adjusting for age and gender (Hare et al., 2020). We first applied the eBass approach and selected the optimal primary threshold by balancing the empirical Bayes 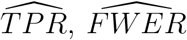 in the objective function (1). The eBass optimal threshold was 0.0018, which was slightly more liberal than *p* < 0.001 (see Figure 3). Next, step two permutation tests were performed based on the primary threshold to identify the significant clusters by controlling FWER < 0.05.

**Figure 3:**
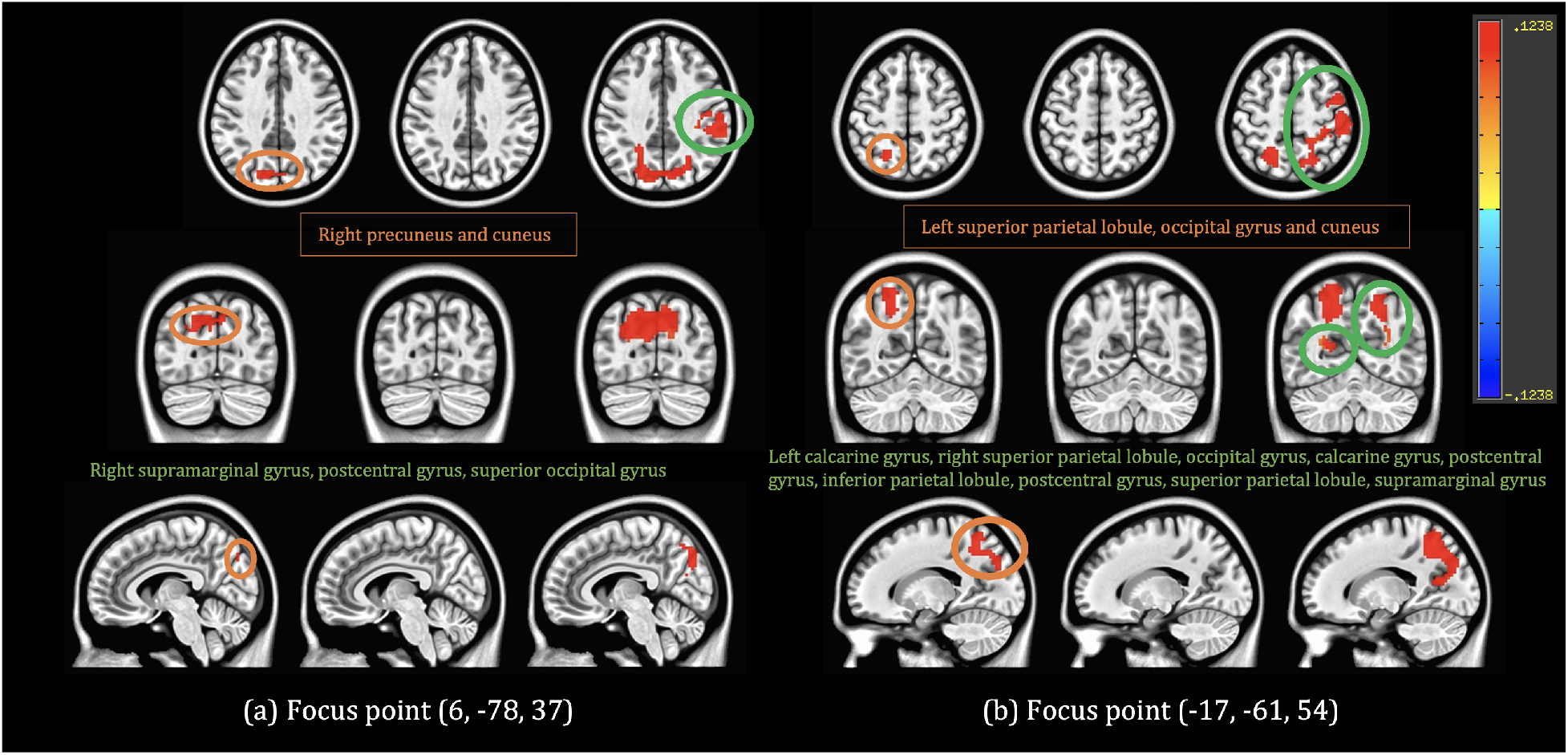
Two groups (in (a), (b)) of significant regions given by Primary thresholds of eBass (column 1), *p* < 0.001 (column 2), and *p* < 0.01 (column 3): axial (row 1), coronal (row 2) and sagittal (row 3) view. (a) focus point is located at (6, -78, 37) on the right brain, (b) focus point is located at (-17, -61, 54) on the left brain. Regions overlap with existing schizophrenia vs nicotine studies were circled by orange line and listed under the first row in the solid frame. The additional regions detected by *p* < 0.01 were circled by green line and listed under the second row without any frame.

We compared the eBass primary threshold with the popular primary threshold values including *p* < 0.001 and *p* < 0.01. We demonstrate the results in Figure 3 as well. All the significant regions were marked in red. The eBass primary threshold based cluster-wise inference yielded significant clusters at left cuneus, superior occipital gyrus and superior parietal lobule, and slightly expanded to right precuneus and cuneus.

In contrast, no significant cluster was detected by the primary threshold of *p* < 0.001 while over-sized clusters were detected by the primary threshold *p* < 0.01 expanding multiple anatomical brain gyri/sulci (e.g., extended to right calcarine gyrus and multiple contiguous regions on right brain, see Fig 3). The clusters based on eBass primary threshold indicate increases in activation at the marked regions, which are highly overlapped with previous findings (Fedota & Stein, 2015; E. Hong et al., 2011). Therefore, the data-driven eBass primary threshold is more flexible and provides a better balance between the sensitivity and false discovery rate, which should lead to greater replicability.

## 5 Discussion

We have developed a data-driven primary threshold selection method for the two-step clusterwise fMRI inference. The multiple comparison problem has been at the heart of neuroimaging data analysis, because it can determine the validity of findings. In practice, true signals in neuroimaging data are often mixed with various sources of noise, and the statistical inference models are sensitive to the noise. Therefore, a small erroneous shift from the optimal decision-making threshold can cause a significant loss of statistical power or uncontrolled false positive findings. However, the primary threshold has been conventionally selected based on empirical analysis and experience, which may not provide the optimal threshold for the target neuroimaging data. To address this need, we propose an empirical Bayes method to calculate estimated sensitivity and false discovery rate and thus facilitate the optimization of selecting the primary threshold for cluster-wise inference.

Built on the successful development of the empirical Bayes approach in the field of high-dimensional statistics, eBass enjoys several advantageous theoretical properties, regarding the estimation robustness and consistency (Efron, 2014; Schwartzman et al., 2009). The eBass threshold provides a reliable cut-off to binarize voxels in the 3D brain space into a point process (Kang et al., 2011). The step two inference (i.e., permutation tests) is also sensitive to the noise level of the point process. When the sensitivity level is low (a stringent threshold), true positive points are unlikely to be spatially adjacent and form a non-trivial cluster resulting in a reduced ability to detect no significant clusters. When a large proportion of false positive points are present in the point process, the false positive points tend to be spatially connected due to the spatial smoothness of the neuroimaging data, leading to cluster-wise false positive findings. For these reasons, we often find it challenging to produce replicable findings in neuroimaging studies (Lindquist, 2020).

Our simulation and data example results concur with the previous findings that the empirical primary threshold (*p* < 0.001) is a good option, especially when no information from the data is available. In general, the primary threshold (*p* < 0.001) can adequately control the false positive findings, which is analogous to the traditional cut of *p* < 0.05 in univariate statistical inference (Eklund et al., 2018). In practice, we find that the data-driven eBass threshold often varies around the primary threshold *p* < 0.001, in many applications. Nevertheless, the eBass primary threshold is objectively selected based on the data, and can thus improve sensitivity in many scenarios (e.g., data sets with smaller sample sizes and small-medium effect sizes). Therefore, we consider the eBass primary threshold to be a good complement to the existing methods for cluster-wise inference. We also note that the eBass is built on the estimation of the two-component mixture model. When the empirical Bayes approach cannot estimate the two components well, we resort to the *p* < 0.001 primary threshold for cluster-wise inference or TFCE as potential solutions.

The eBass method is compatible with all voxel-level statistical inference because the marginal distribution of test statistics is often robust (Chen et al., 2019). The more accurate voxel-level statistical inference can lead to more separable null and non-null distributions and thus more accurate cluster-wise inference results via the eBass primary threshold.

In summary, the eBass provides a data-driven and automatically optimized primary threshold for the two-step cluster-wise fMRI inference. Since the computation is efficient, eBass can be conveniently implemented and compatible with most existing software platforms.

## Appendix

### Theoretical properties of eBass primary threshold

We further provide theoretical results to show the optimality and consistency of the eBass primary threshold estimation.

We first show the eBass solution 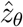 is optimal with respect to the objective function (1). Specifically, we transform the optimization problem in objective function (1) into a saddle point problem (Eban et al., 2017), and solve it by the cost sensitive methods (Parambath et al., 2014).

#### Theorem 1.

*For z_θ_* ∈ Ω_*α*_, *the eBass estimated primary threshold* 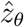 *is optimal with respect to the objective function (1)*.

#### *Proof*.

Following the notation in the method section, we define the decision rule *f_z_θ__*: *f*(*z_v_*) ≥ *f*(*z_θ_*), where *z_θ_* is the primary threshold and *f* is the mixture density. Let *Y*+ be the set of voxels from the non-null component, and *Y*− be the voxels for the null component. Then, let *y_v_* ∈ {−1, 1} be the indicator function where *y_v_* = 1 if *z_v_* ∈ *Y*+ and *y_v_* = −1 if *z_v_* ∈ *Y*−. We consider the 0-1 loss function of true positive and false positives w.r.t. the empirical Bayes estimated 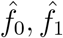. Inherit the notation from Section 2.1, we have

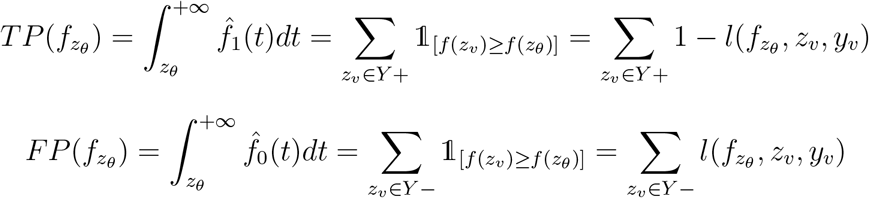

To bound the above two quantities, replacing the 0-1 loss function by the hinge loss function *l_h_*(*f_z_θ__, z_v_, y_v_*) := *max*(0,1 −*y_v_*(*f*(*z_v_*) −*f*(*z_θ_*)) leads to the lower bound of true positives *TP_L_* and upper bound of false positives *FP_U_* at *z_θ_* that satisfies *TP_L_ ≤ TP*(*f_z_θ__*) and *FP_U_ ≥ FP*(*f_z_θ__*) correspondingly.

Once we have the upper bound and lower bound, our objective function 1 can be replaced by a surrogate function 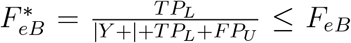. Our goal is to maximize the function 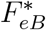 at some given precision. The problem can be written as:

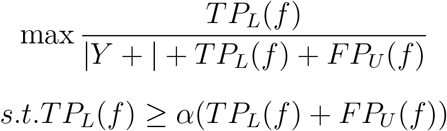

We also replace the notation of the hinge loss function as a shorthand

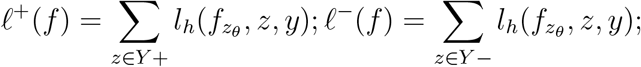

Alternatively, as maximizing 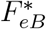 is equivalent to minimizing 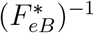, we rewrite the objective function with *ℓ*^+^, *ℓ*^−^:

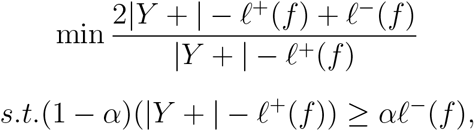

where *α* is given from the 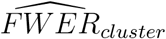 constraint.

Straightforwardly, we can write *ϕ* = |*Y* +| − *ℓ*^+^(*f*), and thus the above minimization is equivalent to

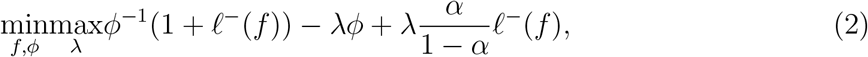

where λ is the Lagrange multiplier. This is similar to optimizing *F_β_* with weighted loss function established by (Eban et al., 2017) where the Cost-Sensitive Classification algorithm was well explained in (Parambath et al., 2014). Since we restrict the search region to the locally convex neighborhood, the solution of this optimization problem is unique. Therefore, our grid search algorithm guarantees the detection of the unique *z_θ_* and achieves the optimality.

Next, we prove the consistency of the eBass estimated primary threshold *z_θ_*. Our proof is mainly built on the fact that the empirical Bayes estimation yields consistent estimators 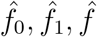 (Petrone et al., 2014; Robbins, 1980).

#### Theorem 2.

*Let* 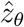 *be the eBass estimated primary threshold, and we have* 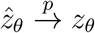.

#### *Proof*.

We let the test statistics of all voxels follow a marginal distribution *z_v_~f*(*z_v_|θ_v_*).

Let ***θ*** = (*θ*_1_, *θ*_2_,⋯, *θ_V_*) be a finite parameter space. Given *θ_ν_* ∈ ***θ***, assume it is independently drawn from a known prior density *π*(***θ***) corresponds to *z_v_*.

Since ***z*** = (*z*_1_,⋯, *z_V_*) is considered a random sample from *f*(·), and the marginal distribution of *f*(**z**) = *∫_**θ**_ π*(*θ*)*f*(**z**|*θ*)*dθ* is proportional to the posterior probability, we can calculate the posterior probability *g*(*θ*|**z**).

As the sequence of estimators in 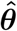 can be estimated by observed *z_v_* with MLE, the empirical Bayes posterior probability is consistent at *θ*_0_ in probability, under the common MLE regularity conditions.

By continuous mapping theorem, our 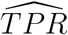 and 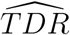 are functions of consistent empirical Bayes estimators. Then we have 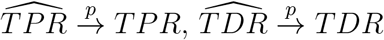. From Theorem 1, *z_θ_* exists and uniquely decided by *TPR* and *TDR*. Therefore, we have 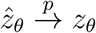.

### Estimating Ω_*α*_ for the step 2 of eBass

In order to control the cluster level FWER below *α*, we should search 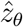 on the set Ω_*α*_. Here, we describe the procedure to identify the set Ω_*α*_ based on the empirical Bayes estimated sensitivity and FDR.

As stated in section 2.2, The cluster level family-wise error rate 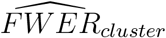 is calculated based on 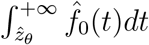. If the total number of false positive voxels 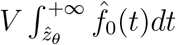 is large at the cut-off of 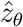, the cluster size of these false positive voxels is likely to be greater than the permutation tests determined cluster-size threshold *K_α_*. In results, false positive clusters appear in the final results. In order to avoid the false positive cluster, the cut-off 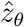 is required to prohibit forming large clusters.

Specifically, we denote the estimated number of false positive voxels using a cut-off 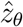 by 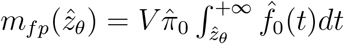. We next compute the upper bound of the cluster size based on the combinatorial probability that these 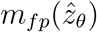 false positive supra-threshold voxels can form a contiguous non-trivial cluster in the three-dimensional brain space. We define the search domain for 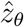 by Ω_*α*_ = {*z_θ_*: Sup{*μ*(*m_fp_*(*z_θ_*))} < *K_α_*}, where *μ* is the cardinality measure of any set of contiguous voxels formed by *m_fp_*(*z_θ_*) in the brain space. In practice, the direct calculation of Sup{*μ*(*m_fp_*(*z_θ_*))} is intractable. We resort to permutation based techniques to approximate Sup{*μ*(*m_fp_*(*z_θ_*))}.

In each permutation, the random shuffling of subject labels hypothetically produces a 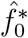 distribution and an *α_p_* level is chosen to control the permutation test FWER. Since we are able to theoretically calculate the 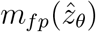 based on empirical Bayes estimated 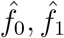, we consider 1) an *α*_1_ for permutation bound that controls the FWER among all supra-threshold voxels, 2) an *α*_2_ for false positive cluster bound controls the FWER for estiamted false positive voxels. Commonly, the widely used 5% *α_p_* level is unadjusted and that *α_p_ = α*_1_. The adjustment according based on the definition of *α*_1_ and *α*_2_ is (1 −*α*_1_)(1 −*α*_2_) = 1 −*α_p_*. For *α*_2_, it is calculated by the top 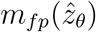 voxels on both tail-end of 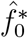 and randomly choose 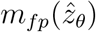 from those extreme observations to form false discovery clusters. The adjustment indicates that when *α*_2_ → 0, 1 − *α*_1_ ≈ 1 − *α_p_*. In other words, we need to take the cluster size corresponds to *max*{*α*_2_} (or other levels based on the adjustment) to estimate 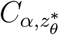, and then we can estimate the support Ω_*α*_.

We summarize the procedure as follows:

Step 1: For a cut-off 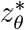, we calculate 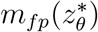, the number of false discoveries based on the empirical Bayes estimated parameters.
Step 2: Compute *K_α_* by shuffling subject labels at the cut-off 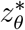.
Step 3: Shuffle the subject labels for *J* times. At each iteration *j* we select 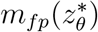 voxels with the largest test statistics and a similar dependence structure.
Step 4: Let 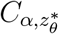 denote the maximum (or other adjusted α level) cluster size across the J permutations in step 3.
Step 5: 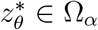 if 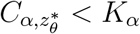.

In practice, we find both 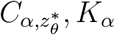 are monotonously decreasing with 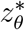 while the decreasing speed of 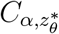 is faster. Therefore, Ω_*α*_ is often a continuous domain.

### Comparison between Cluster-extent Methods and Voxel-wise Inference

Although we focus on the two-step cluster-wise inference in the simulation study, we further compare the cluster-extent methods with TFCE - one of the most popular voxel-extent inference. The TFCE generates voxel-level corrected *p* values instead of considering clusters as a whole, thus the cluster-wise FWER is not applicable to this method. Similarly, as in cluster-wise inference we assume all supra-threshold voxel-formed clusters are significant, we compare the voxel-wise sensitivity and FDR of primary thresholding step specifically with TFCE.

The 2D image has the same dimension with *V* = 100 × 100 = 10, 000 voxels. The truth consists of four identical squared areas that are placed in the center of the image with equal distance (roughly same as the side of the square). Different from the images in the simulation section, we apply the smoothing step after adding the true signals to the original image. In this way, the underlying truth have an irregular shape on the margin, and the strength of signal decreases steady from center to margin. The total number of true significant voxels is 900. We perform the simulation study on the set of images have ES=0.6, smoothed with FWHM=8mm, 30 subjects per group.

We perform the two-step cluster-wise inference with eBass, BH-FDR correction, *p* < 0.001, and *p* < 0.01. We also apply TFCE voxel-wise inference on the images under this setting. Similarly, we evaluate the performance of methods by their voxel-wise sensitivity, FDR, and cluster-wise FWER. We exhibit the simulation results in Table 2.

**Table 2.** Performance comparison between cluster-extent methods and voxel-wise thresholding method. 30 subjects per arm with ES=0.6, smooothed with Gaussian kernel FWHM=8mm.

|  | eBass | BH-FDR | <0.001 | <0.01 | TFCE |
| --- | --- | --- | --- | --- | --- |
| Primary Threshold | 0.003±0.0029 | 0.0063±0.0001 | 0.001 | 0.01 | NA |
| Sensitivity | 0.9996±0.0007 | 0.9998±0.0004 | 0.9991±0.001 | 0.9998±0.0004 | 1 |
| vFDR | 0.2469±0.025 | 0.2644±0.0129 | 0.2200±0.0096 | 0.2767±0.0116 | 0.3602±0.0142 |
| cFWER | 0 | 0 | 0 | 0 | NA |

From the example above, we find out that when the significant clusters have blurred edges and close to each other, the voxel-extent methods would have a significant increase (up to 50%) in vFDR comparing to cluster-extent methods. When the ES and sample size are low to medium, the voxel-extent method (e.g., TFCE) tends to outperform cluster-extent methods on sensitivity by 5-30% in this type of images who have clear edges, while the vFDR remains about the same.

### Primary Thresholds for Real Data

**Table 3.** Primary thresholds and corresponding cluster-extent thresholds

| eBass | Hard Threshold p<0.001 | Hard Threshold p<0.01 |
| --- | --- | --- |
| 0.0018(≈154) | 0.001(105) | 0.01(853) |

